# A Label-free Nanowell-based Impedance Sensor for Ten-minute SARS-CoV-2 Detection

**DOI:** 10.1101/2024.12.11.627986

**Authors:** Zhuolun Meng, Liam White, Pengfei Xie, S. Reza Mahmoodi, Aris Karapiperis, Hao Lin, German Drazer, Mehdi Javanmard, Edward P. DeMauro

## Abstract

This work explores label-free biosensing as an effective method for biomolecular analysis, ensuring the preservation of native conformation and biological activity. The focus is on a novel electronic biosensing platform utilizing micro-fabricated nanowell-based impedance sensors, offering rapid, point-of-care diagnosis for SARS-CoV-2 (COVID-19) detection. The nanowell sensor, constructed on a silica substrate through a series of microfabrication processes including deposition, patterning, and etching, features a 5*×*5 well array functionalized with antibodies. Real-time impedance changes within the nanowell array enable diagnostic results within ten minutes using small sample volumes (<5 µL). The research includes assays for SARS-CoV-2 spike proteins in Phosphate-buffered saline (PBS) and artificial saliva buffers to mimic real human SARS-CoV-2 samples, covering a wide range of concentrations. The sensor exhibits a detection limit of 0.2 ng/mL (1.5 pM) for spike proteins. Middle East Respiratory Syndrome (MERS-CoV) spike proteins are differentiated from SARS-CoV-2 spike proteins, demonstrating specificity.

**TOC Graphic:** 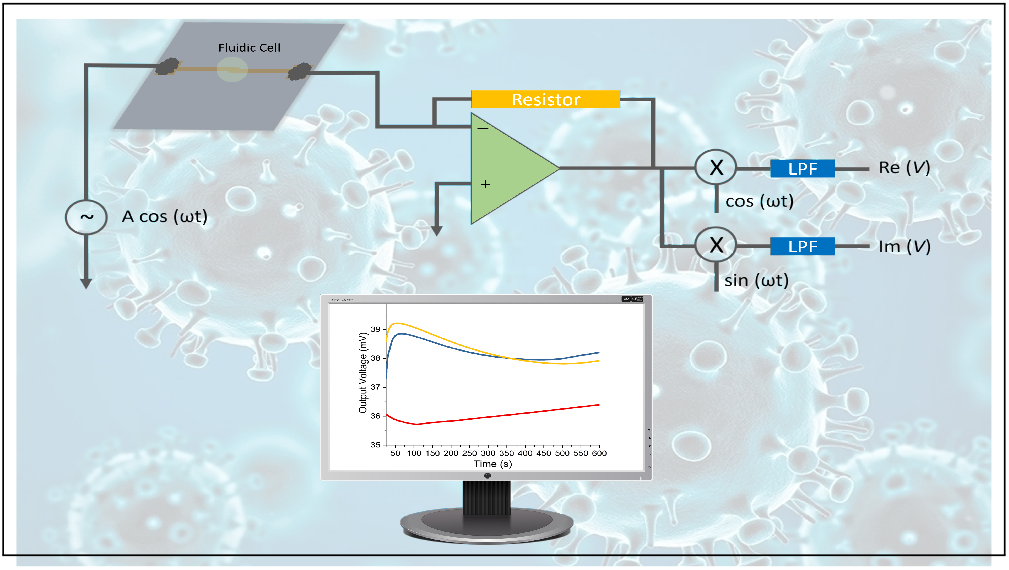

## INTRODUCTION

Coronavirus Disease 2019 (COVID-19) is an infectious contagious disease caused by the SARS-CoV-2 virus. In December 2019, the first known case of COVID-19 was identified in Wuhan, China.^1^ In the following months, the SARS-CoV-2 virus rapidly spread across the world and now has multiple mutations.^2–4^ According to the World Health Organization (WHO) data, the cumulative total reported cases by May 12, 2024, was 775,481,326.^5^ COVID-19 significantly impacted the global economy, food security, education, and mental health, among other effects.^6^

There are ongoing efforts to fight against respiratory diseases with high transmissibility by researchers and scientists from different disciplines. ^4,6–8^ The development of vaccines and treatment strategies has successfully decreased hospitalization and mortality rates. ^9–11^ In addition, to potentially control the spread of the disease, detection is the first line of defense and is one of the successful responses to the pandemic.^8,12^ Testing is also pivotal for public health, and detection of SARS-CoV-2 helps investigators characterize its prevalence, spread, and contagiousness. ^8^

There are multiple diagnostics methods developed for COVID-19 in the past few years, which can be categorized into RNA & DNA/ molecular diagnosis, antibody/ antigen testing, clinical imaging techniques, and biosensors.^13–16^ RNA & DNA/ molecular diagnosis are some of the most developed detection methods. ^14,15^ They are highly sensitive, accurate, and specific for SARS-CoV-2 virus detection. ^14,15^ However, these methods have disadvantages, including the need for trained operators and long workflow times (from 30 minutes to several days).^14,15^ Antibody/ antigen testing methods have specific advantages, including large capacity, rapid results, inexpensive materials, portability, and ease of operation.^13–15^ These methods, however, are not as accurate as molecular diagnosis.^13–15^ Other detection methods are based on medical imaging techniques, especially computed tomography (CT), X-ray imaging, and ultrasound, which analyze chest images to diagnose patients. These detection methods are non-invasive and could be implemented for fast screening, especially in combination with automated image analysis.^14,15,17^ However, the equipment cost and radiation exposure need to be taken into consideration, particularly in the case of CT scans, which expose patients to significant amounts of radiation and cannot be used frequently.^14,15,17^ As the technologies develop, biosensors are becoming a reliable option for disease detection and diagnosis. Compared to the detection methods discussed above, biosensor-based methods present alternatives that do not need advanced equipment and skilled operators for rapid diagnosis.^15,18–21^ In particular, label-free devices for bio-detection have developed significantly over the last decade. Detection utilizing label-free devices of biomarkers has numerous advantages compared to label-based counterparts, including cost-effectiveness, simpler sample preparation, a broad range of targets, portability, and Point-of-Care capabilities. ^22–24^

In this study, a microfabricated label-free nanowell array impedance sensor is used to detect SARS-CoV-2 spike proteins in artificial saliva. In previous research, this sensor was used to detect stress hormones and cytokine in serum.^25–30^ A new preparation and data analysis method for a nanowell sensor is presented which demonstrates a lower limit of detection (LOD). Additionally, the sensor can discriminate between SARS-CoV-2 spike proteins and MERS-CoV spike proteins, demonstrating target specificity.

## MATERIALS AND METHODOLOGY

### Impedance Sensor Methodology

The sensor is a 5*×*5 array of wells microfabricated over a 20 µm *×* 20 µm area consisting of two opposing gold electrodes separated by an aluminum oxide layer. Antibodies are first injected and adsorbed in the wells. A sample is then introduced and the impedance between electrodes is monitored through lock-in amplifiers to determine the presence of the corresponding antigen. The schematic cross-section view of a single well in the array is shown in Fig. 1a, indicating the two gold layers acting as electrodes and separated by a thin dielectric layer of aluminum oxide. The equivalent circuit model is shown in Fig. 1b, and discussed in detail in previous research by Mahmoodi et al.^27^ The following is a brief description of the testing procedure. First, PBS buffer is injected inside the wells to create a liquid environment. Then, the selected antibodies are injected into the wells and adsorbed on their surface. The changes in impedance between the two gold electrodes are monitored in real-time using a lock-in amplifier to determine if the antibodies adsorbed successfully, as shown schematically in Fig. 1c (see a top view schematic in Figure S1a in the supplementary material). Subsequently, the test sample is introduced into the wells, and the changes in impedance are continuously monitored to detect the binding of antigens onto the antibodies (see schematic of the binding in Fig. 1d and the corresponding top view schematic in Fig. S1b in the supplementary material). Measurable increases in impedance indicate the presence of antigen-antibody pairs after introducing the test sample.

**Figure 1:**
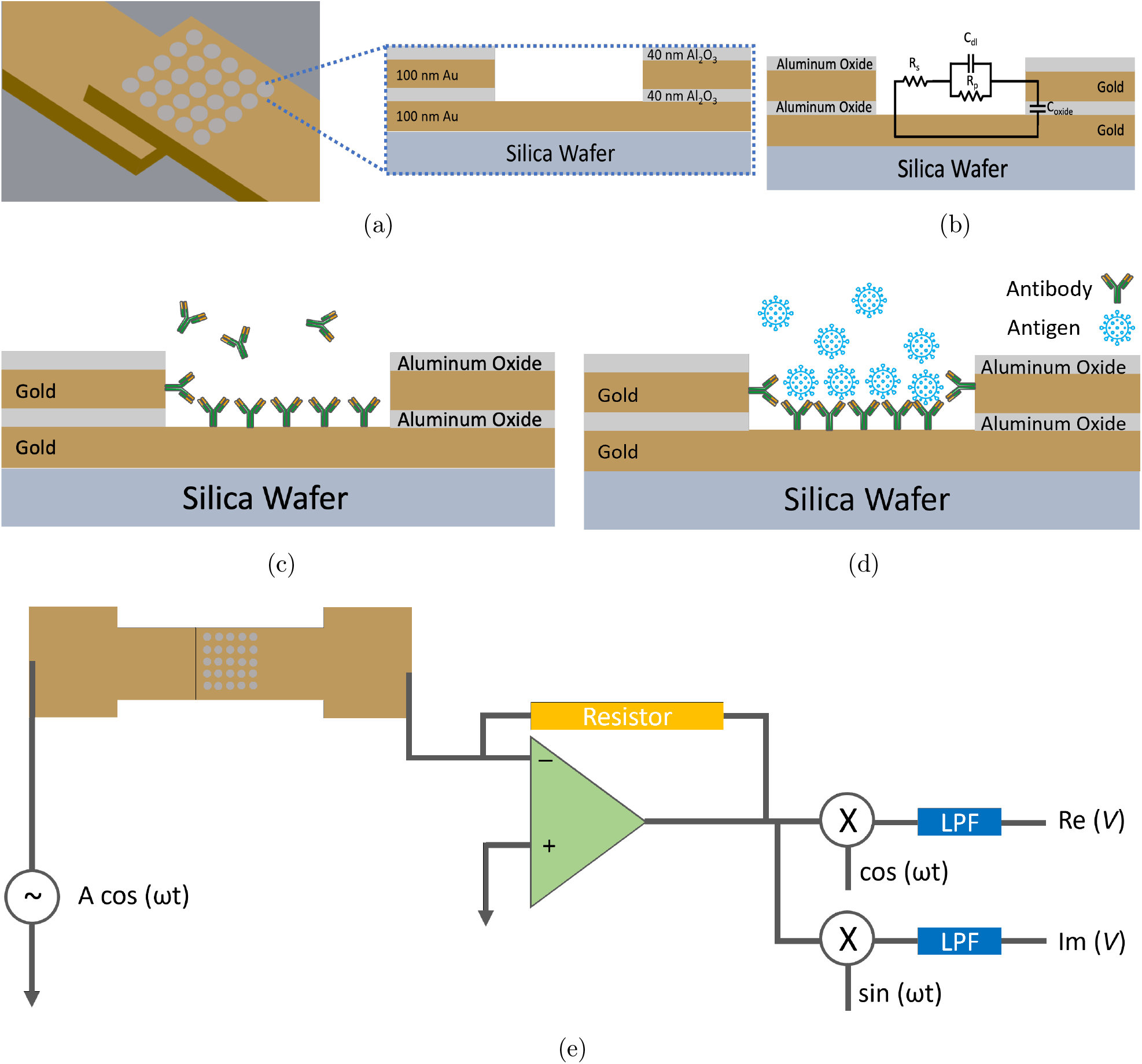
(a) Cross-sectional view of a single nanowell. (b) Equivalent circuit model. (c) Cross-sectional view of single nanowell adsorbing antibodies. (d) Cross-sectional view of the nanowell adsorbing target proteins. (e) Equivalent circuit of measurement platform using a lock-in amplifier.

### Sensor Fabrication

A schematic of the key fabrication steps to create the nanowell sensor is presented in Fig. 2a. The sensor is prepared on 7.62 cm-diameter, 500 µm-thick fused silica substrates. The first layer of gold (100 nm) is deposited by physical vapor evaporation (E-beam) on a silica wafer previously covered with a 5 nm chromium layer to improve adhesion. The first electrode is then created from this gold layer by photolithography and lift-off processing (see first step, Fig. 2a). A 40 nm layer of aluminum oxide is subsequently deposited by atomic layer deposition (second step, Fig. 2a). A second 5 nm chromium adhesion is then deposited, followed by a second 100 nm gold layer deposition using the same process as the first layer (third step, Fig. 2a). Note that the first and second layers of gold overlap in a small 20 µm *×* 20 µm area. Lastly, a 40 nm aluminum oxide layer is deposited as a protection layer on top (fourth step, Fig. 2a). After depositing all metal layers, multiple wet etching steps are performed to pattern the well-shaped array on the overlapping area between the two gold electrodes by coating a layer of photoresist (PR) and etching the two aluminum oxide layers (buffered oxide etchant), one gold layer, and one chromium layer (gold and chromium etchants) to expose the bottom gold layer (see zoom-in view of the array in Fig. 2a, fifth step). After the strip-off of the PR, a second mask is used to feature another layer of PR to etch the aluminum oxide outside the sensor to remove the residue on the silica substrate. A polydimethylsiloxane (PDMS, Sylgard 184, Dow Corning, 10:1 prepolymer/curing agent) fluidic cell is glued (5 Minute Epoxy, Devcon, Illinois Tool Works Inc.) above the array of wells to keep the liquids contained in the working area (shown in Fig. 2a, sixth step). To connect the impedance sensor to other electronic equipment, electrically conductive wires are attached to the gold connection pads by conductive epoxy (CW 2400, Chemtronics, Kennesaw, GA, USA), as shown in the last step in Fig. 2a. In Fig. 2b are views of the nanowell array with different magnification, including a view of the wafer after fabrication, a single sensor, the nanowell structure, and the array of wells observed under a bright-field microscope (Ernst Leitz GmbH, Wetzlar, Germany).

**Figure 2:**
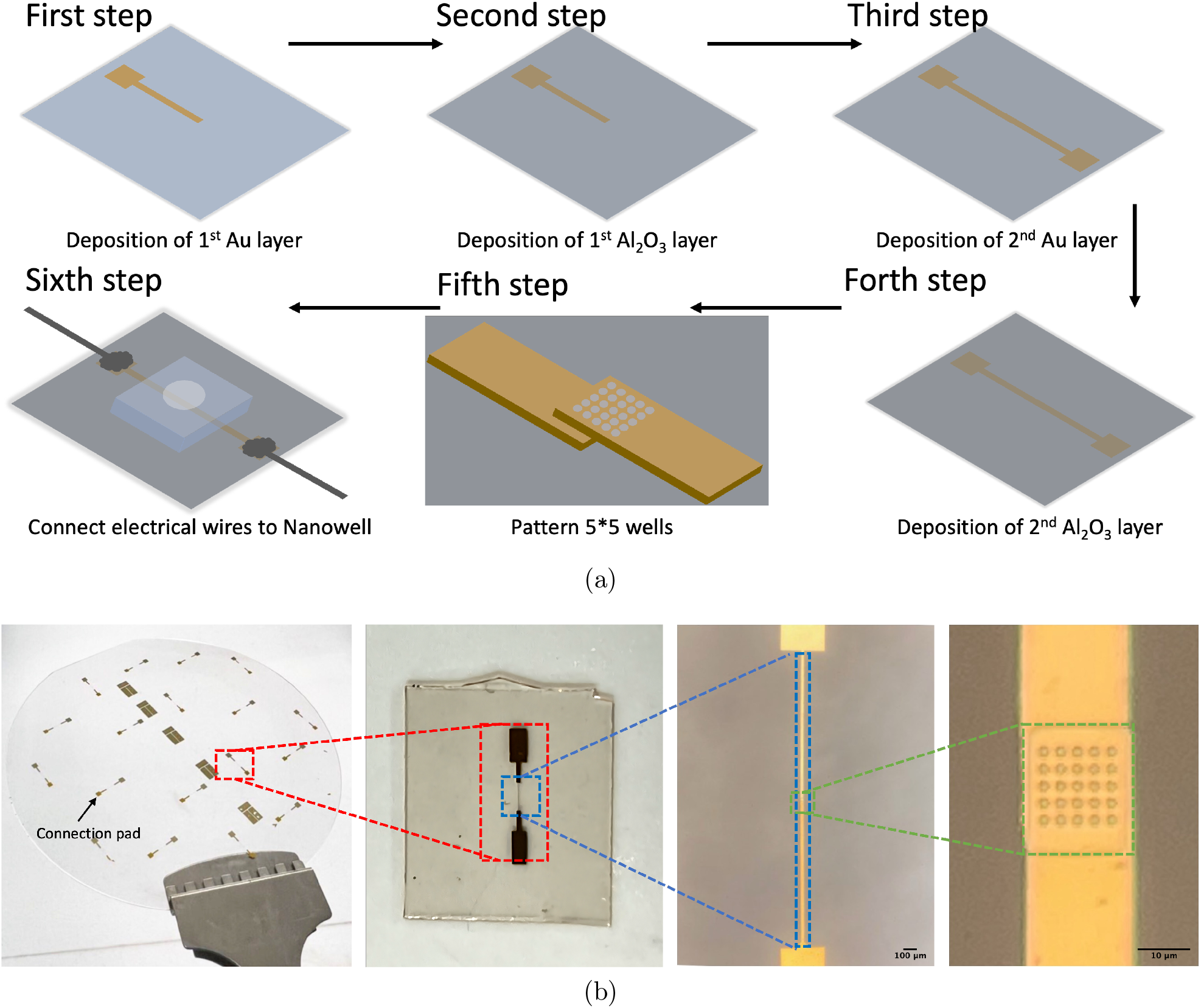
(a) Microfabrication procedures for nanowell sensors. Follow the arrows: First step: First 100 nm of the gold electrode after deposition and lift-off; Second step: First 40 nm *Al*_2_*O*_3_ layer by ALD; Third step: Second 100 nm gold electrode using the same fabrication processes as the first layer; Fourth step: Second 40 nm *Al*_2_*O*_3_ layer by ALD; Fifth step: Zoom-in view of Well-shaped arrays in the middle of overlapping area are exposed by multiple etchings for two *Al*_2_*O*_3_ layers and one gold layer; Sixth step: Finish connection setup with conductive wires and epoxy, and install PDMS fluidic cell. (b) From left to right:1. View of a fabricated wafer; 2. Single nanowell sensor; 3. Microscope view of electrodes; Microscope view of 5 *×* 5 well-shaped arrays.

### Preparation of the Reagents

Polyclonal and monoclonal SARS-CoV-2 antibodies (R&D Systems, Minneapolis, MN, USA) are used throughout the experiments in a 100 µg/mL concentration in PBS. No significant differences in results are seen between the type of antibody used and Fig. S4 shows the overlapping responses of the monoclonal and polyclonal antibodies. The first experiments are performed using SARS-CoV-2 spike protein (R&D Systems, Minneapolis, MN, USA) as the target protein and are prepared in artificial human saliva (Sigma-Alrich, St. Louis, MO, USA) in concentrations ranging from 0 – 1000 ng/mL (0 - 7.5 nM). In addition, MERS-CoV spike protein (University of California-San Diego, San Diego, CA, USA) is used to perform a specificity test on a different protein, prepared within 0.18X PBS at a concentration of 1000 ng/mL. The PBS used in this work is 1X (1X PBS, pH=7.4, Gibco, Thermo Fisher Scientific Inc., Waltham, Massachusetts, US). The diluted PBS used later in this work is diluted by deionized (DI) water from Rutgers Micro Electronics Research Laboratory.

### Real-time Measurements

Fig. S2 shows the real-time impedance measurement for the first pipetting of (5 µL) 1X PBS to the empty nanowell sensor, monitored under 4 different frequencies. For 1X PBS, following an initial increase caused by pipetting PBS into an empty sensor, which increases the media’s voltage baseline, the output voltage subsequently decreases, approaching a constant voltage level. The voltage gradually increases without the existence of any adsorbable material, possibly because of the slight and persistent evaporation inside the sensor. Increasing the frequency results in a higher impedance value. For example, in Fig. S2, curves from top to bottom represent frequencies from 2 MHz to 100 kHz. 1 MHz is used during the experiments to avoid the inductive effect in the system at high frequencies.^26–28^

The experiments presented in this work have the following procedure. First, the sensor is prepared by connecting to the lock-in amplifier and is supplied with a 100 mV, 1 MHz AC signal. The first solution added to the sensor is 5 µL of 1X PBS, and for all steps, the solution resides in the sensor for 10 minutes, undisturbed, before the subsequent step commences. Next, 3 µL more of PBS are added, followed by 3 µL of antibodies prepared in PBS that are adsorbed to the surface of the sensor. The power source is then removed, and the 11 µL of liquid are removed from the sensor via a pipette. After the liquid is fully removed, the power is restored, and two more rounds of PBS are added, as mentioned above, to recreate the liquid environment. Lastly, 3 µL of SARS-CoV-2 spike protein prepared in artificial saliva of the target concentration is added to the sensor.

Fig. 3A is a plot of the recorded voltage outputs. Shown in red is the voltage response of the first round of 3 µL of PBS added to the sensor, blue is the addition of the antibody solution, and yellow is 1000 ng/mL antigen suspended in artificial saliva. Before the real-time test, the sensor is prepared with antibodies. In this figure, the blue curve represents the impedance change during antibody injection. As the curve descends, it indicates that the antibodies are binding to the sensor, which increases impedance and causes a decrease in voltage. When the liquid solutions are added, inserting the pipette tip into the well causes large shifts in output voltage before stabilizing over a short period of time. The red PBS curve displays a slowly increasing voltage due to the enhanced conductivity provided by the PBS. The addition of the antibodies results in a decrease in voltage due to their adsorption to the sensor surface and demonstrates that the sensor is functional. Fig. 3b is an isolated view of the yellow spike protein response curve. Due to every experiment having slightly different pipette insertion times after the recording is started, three time instances: *t*_0_, *t*_*ref*_, and *t*_*end*_ are used to calculate the decrease in voltage for each trial and will be used to evaluate the performance of the sensor. *t*_0_ is the location of the last shift in voltage due to pipette insertion, *t*_*ref*_ is 30s after *t*_0_, and *t*_*end*_ is 580s after *t*_0_. The decrease in voltage is then calculated as 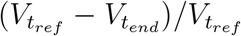 A previous method to calculate the voltage drop is to instead use the time of the relative maximum after *t*_0_ as *t*_*ref*_. ^25,27,28^ Both methods will be used and compared in this study. Using the first method results in a voltage decrease of 3.64%, shown in green, and the second is 3.70%, shown in red. The new method was developed due to some signals not having a clearly defined relative maximum. Thus, the new method is a more robust approach to interpreting the results.

**Figure 3:**
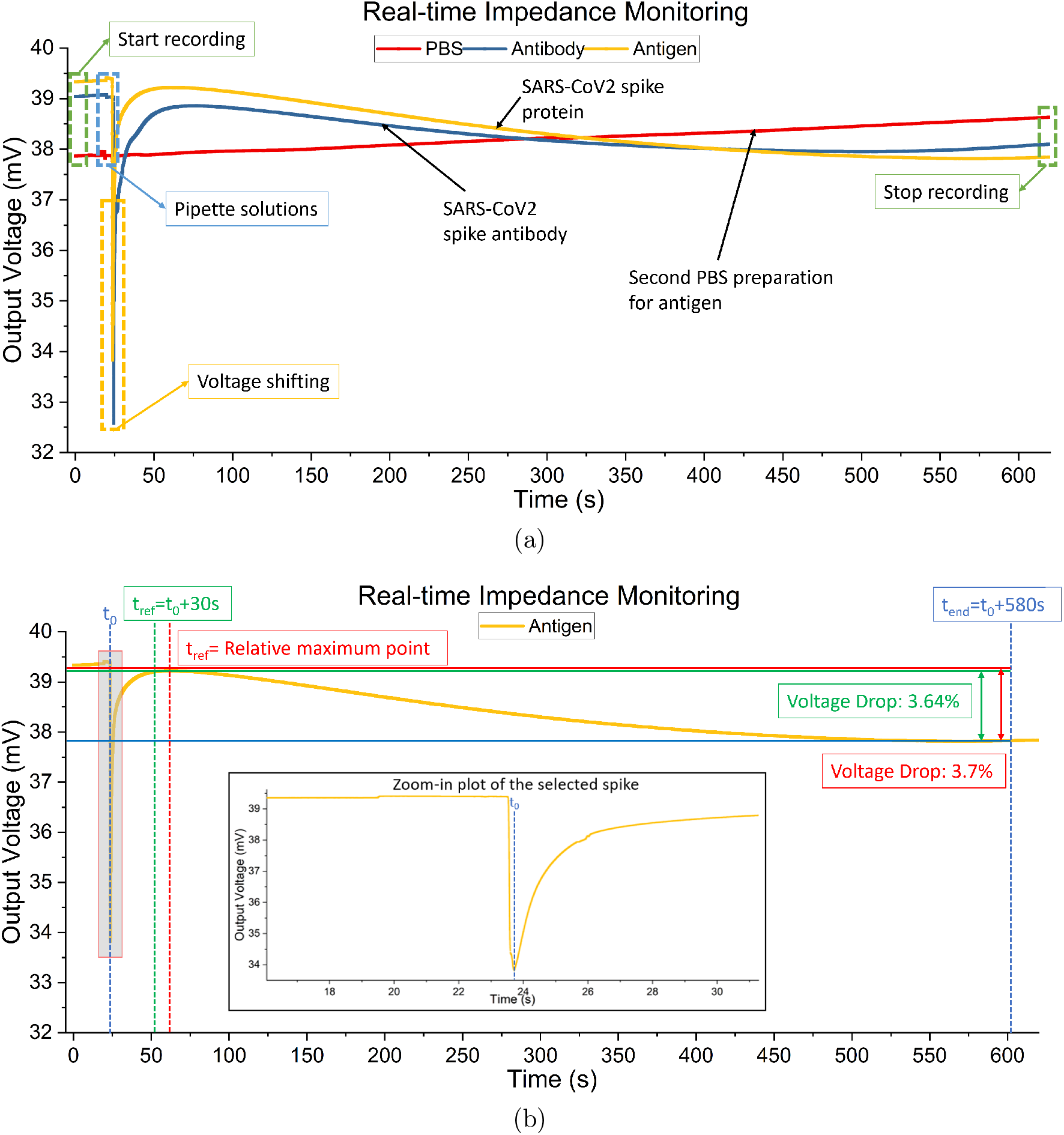
An example of the data analysis methods using 1000 ng/mL SARS-CoV2 spike protein samples. (a) Comparison of real-time voltage monitoring after pipetting 1X PBS, antibodies, and antigens. (b) Comparison of two analysis methods.

## RESULTS AND DISCUSSION

### SARS-CoV-2 Spike Protein Responses in Artificial Saliva

Experiments are performed using antibodies in 1X and 0.18X PBS solutions paired with SARS-CoV-2 spike proteins suspended in artificial saliva. The first experiments use antibodies in 1X PBS solutions. Fig. 4a shows the data analysis results based on the first method described above, while Fig. 4b demonstrates the analysis results using the second method.^25,27,28^ From these two figures, the negative control (NC) (C = 0 ng/mL) overlaps with 100 - 200 ng/mL. Thus, the LOD is estimated to be not less than 200 ng/mL (1.5 nM, molar concentration = (mass of solute/volume of solution)*×*(1/molecular weight) where the molecular weight is 134 kDa for SARS-CoV-2 Spike Protein) which is not ideal and could be due to the differences in impedance between the 1X PBS and artificial saliva.

**Figure 4:**
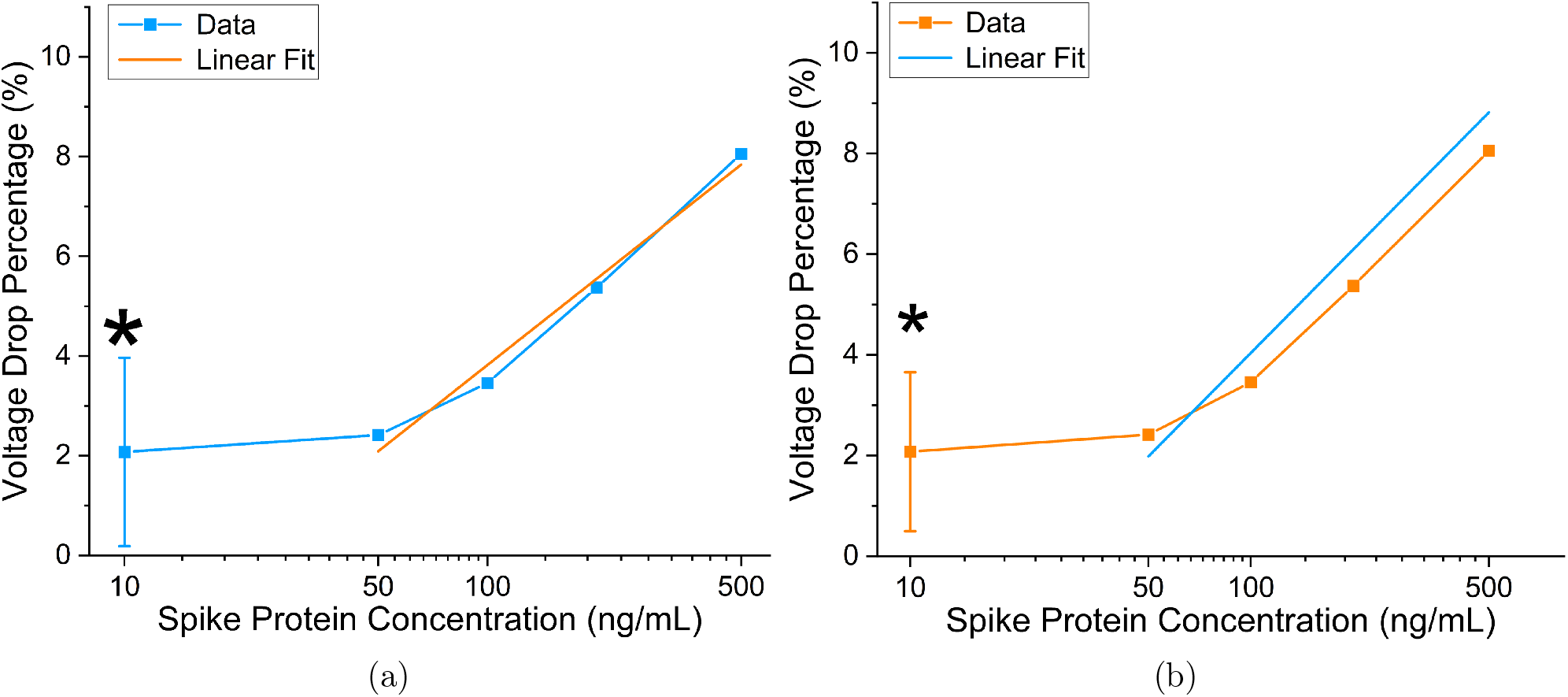
Sensing of spike protein in saliva with antibody in 1X PBS solution. (a) Titration curve using updated data analysis method for SARS-CoV-2 spike proteins in saliva and antibodies in 1X PBS with 50-500 ng/mL dynamic range and 200 ng/mL (1.5 nM) detection limit; linear fit: (V) = −7.68 + 5.75 *×* (C), R-Square = 0.98. (b) Titration curve using previous data analysis method for SARS-CoV-2 spike proteins in saliva and antibodies in 1X PBS with 50-500 ng/mL dynamic range and 200 ng/mL (1.5 nM) detection limit; linear fit: (V) = −9.64 + 6.84 *×* (C), R-Square = 0.96. ⋆ Denotes negative control (C = 0ng/mL) plotted at C = 10 ng/mL for visualization purposes.

As we speculate the impedance difference between 1X PBS solution and saliva may negatively affect detection sensitivity, we next seek to use a PBS solution that has matching impedance. Fig. S3a shows the voltage responses for five different concentrations of PBS, ranging from 0.1X to 1X, and artificial saliva. In Fig. S3b, the output voltage for artificial saliva is between 0.1X and 0.2X PBS. A regression curve is then plotted for the different PBS solutions to find the equivalent PBS concentration for saliva. The output voltages at 300 seconds are used for all tests, and the equivalent PBS concentration for saliva is found to be 0.18X PBS through interpolation.

Fig. 5a and 5b show the titration curves for antibodies suspended in 0.18X PBS buffer to match the baseline of saliva. Fig. 5a uses the new data analysis method, and Fig. 5b uses the previous method. For these results, concentrations were titrated from 0.1 ng/mL - 1000 ng/mL. Fig. 5a displays a statistically significant difference in voltage response between 0 and 0.2ng/mL (1.5pM) using a 0.05 significance level, which is a three-order-of-magnitude improvement compared to the estimated limit of detection using 1X PBS. When using an even higher significance level of 0.001, the detection limit is 0.5ng/mL (3.7pM) and is still substantially higher than the previous estimate. Fig. 5b displays a 1ng/mL (7.5 pM) limit of detection, using a 0.001 significance level and the new data analysis method. This result, while worse than the new analysis method, is also an improvement over the previous detection limit. The detection limits for significance levels from 0.05 to 0.0001 can be seen in Table S1. The linear fits can be seen in both plots, and the intersection of the fits and the negative controls result in theoretical detection limits of 0.13ng/mL (0.975pM) and 0.33ng/mL (2.475pM) for the new and previous methods, respectively.

**Figure 5:**
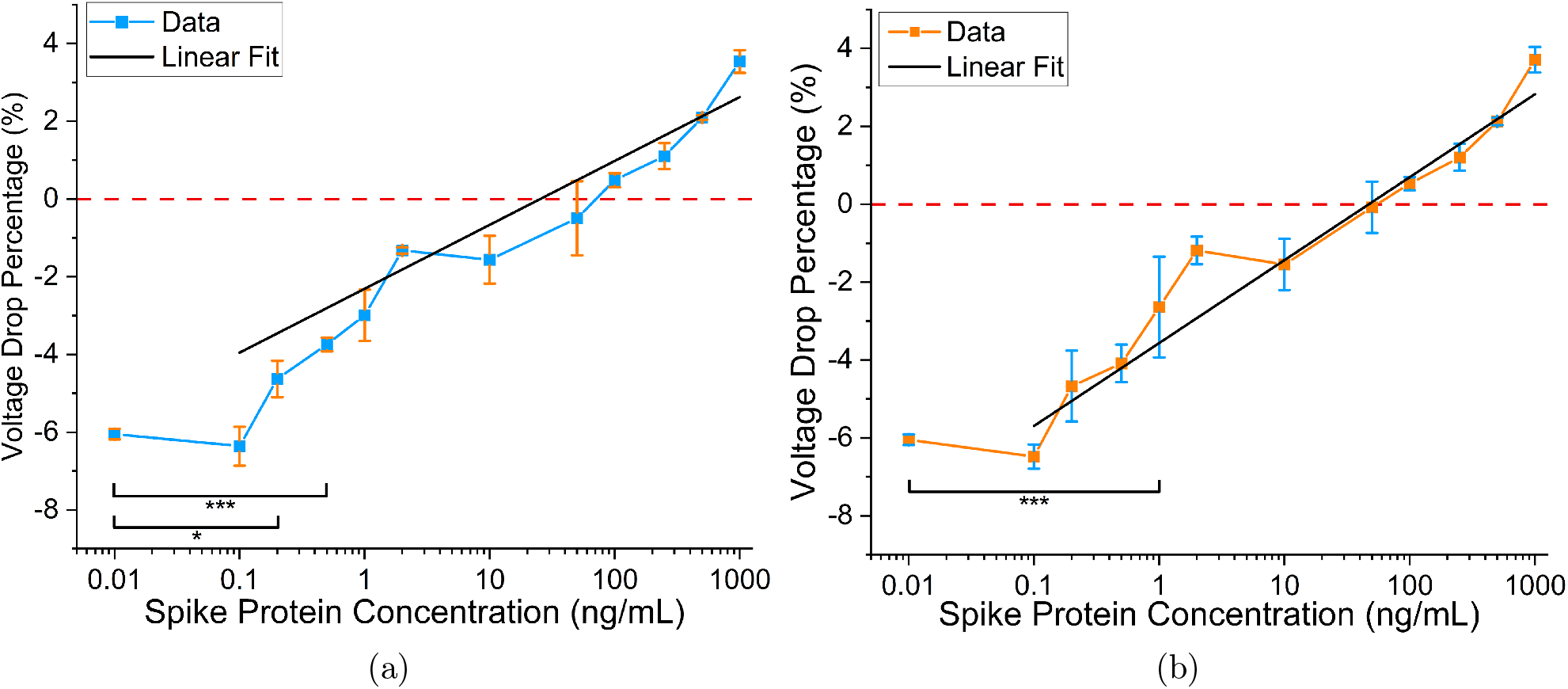
Sensing of spike protein in saliva with antibody in 0.18X PBS solution. Negative control (C = 0ng/mL) plotted at C = 0.01 ng/mL for visualization purposes. (a) Data using updated data analysis method for SARS-CoV-2 spike proteins in saliva and antibodies in 0.18X PBS with 0.1-1000 ng/mL dynamic range; linear fit: (V) = −2.3 + 1.64 *×* log (C), R-Square = 0.95. (b) Data using previous data analysis method for SARS-CoV-2 spike proteins in saliva and antibodies in 0.18X PBS with 0.1-1000 ng/mL dynamic range; linear fit: (V) = −3.56 + 2.13 *×* log (C), R-Square = 0.96. ⋆*P ≤* 0.05; ⋆ ⋆ ⋆*P ≤* 0.001

The LODs for antibodies prepared in 0.18X PBS solutions are much lower than those in 1X PBS solutions. All titration curves in Fig. 4 and 5 show similar and strong linear relationships; however, Fig. 5 displays much lower LODs than Fig 4. Using 0.18x PBS, the new and old analysis methods result in experimental LODs of 0.2 (1.5 pM) and 1 ng/mL (7.5 pM) respectively, compared to 200 ng/mL (1.5 nM) with the 1x PBS solution. Thus, 0.18X PBS is a more suitable buffer for SARS-CoV-2 spike antibodies for this application. Lastly, the agreement between the two data analysis methods for the antibodies prepared in 1X and 0.18X PBS gives credence to the matched baseline voltage having a significant impact on the LOD and is not artificially lowered by the data analysis method employed.

### MERS-CoV Specificity Tests

After demonstrating that the sensor can detect a binding event between SARS-CoV-2 spike protein and a matching antibody, the specificity of the sensor is examined by applying MERS-CoV spike proteins to a sensor prepared with SARS-CoV-2 antibodies. MERS-CoV is used for the specificity test as it is a coronavirus closely related to SARS-CoV-2.^31^ In these experiments, MERS-CoV spike protein with a concentration of 1000 ng/mL in 0.18X PBS is introduced to the sensor using the same procedure as before. In previous experiments, there are no obvious differences between using artificial saliva and 0.18X PBS as protein buffers. A spike protein concentration of 100 ng/mL prepared in artificial saliva and 0.18X PBS had voltage drops of 0.49 and 0.64% respectively, as shown in Fig. S5, which were within uncertainty bounds. Therefore, the use of a 0.18x PBS buffer is not expected to affect the specificity tests for MERS-CoV spike protein in this section.

Fig. 6 shows the results of the specificity tests and displays that the sensor can differentiate between the two spike proteins. Two negative controls (Negative Control 1 and Negative Control 2 in Fig. 5) tests are shown in blue and green, two MERS-CoV spike protein curves result in black and orange, and a SARS-CoV-2 spike protein results in red (250 ng/mL). All curves are normalized at 50 seconds to reduce noise from the sensor’s voltage baseline, allowing for a clearer comparison. The MERS-CoV spike protein responses overlap with the negative controls, demonstrating that the sensor does not detect a binding event between the MERS-CoV spike protein and SARS-CoV-2 antibody and instead increases in voltage due to the presence of the PBS buffer, thus displaying the ability of the sensor to differentiate between the two antigens.

**Figure 6:**
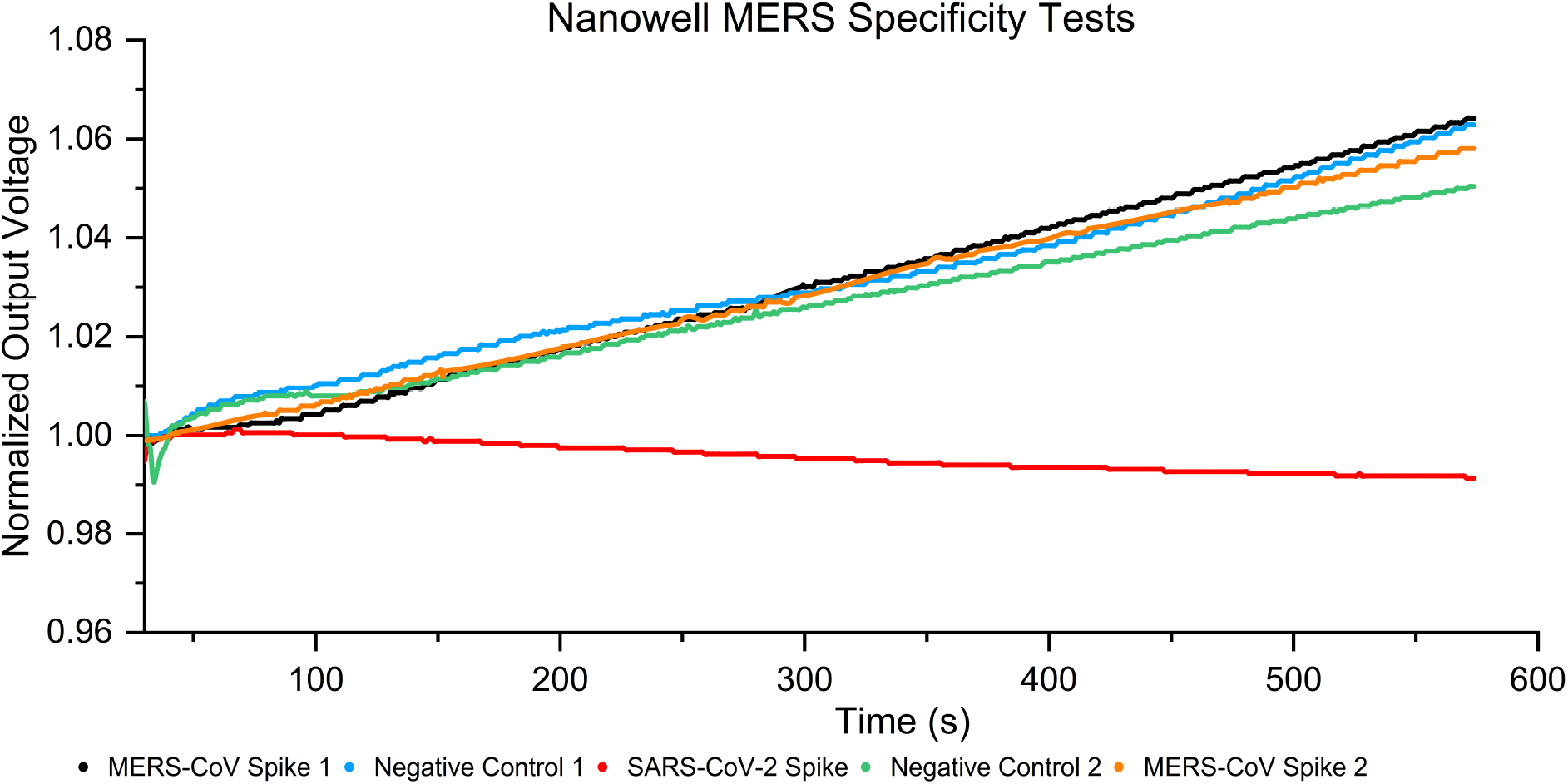
Comparisons between MERS-CoV spike proteins, SARS-CoV-2 spike proteins, and negative control using SARS-CoV-2 spike antibodies. Only the red SARS-CoV-2 spike protein curve gives a response to the SARS-CoV-2 antibodies, and MERS-CoV curves follow the same trends as the negative control curves, proving the device is capable of differentiating SARS-CoV-2 proteins from similar respiratory disease samples.

## Conclusions

A label-free and rapid microfabricated impedance biosensor is presented that can detect SARS-CoV-2 spike protein successfully. SARS-CoV-2 spike proteins in saliva with antibodies in 1X PBS displayed an experimental limit of detection of 200 ng/mL (1.5 nM) within 10 minutes. To lower the LOD, the baseline voltage output of different concentrations of PBS was investigated to find the best match for the artificial saliva. Through a regression curve, 0.18X PBS was the resultant match, and the experimental LOD with this concentration of PBS was lowered to 0.2 ng/mL (1.5 pM). Apart from the comparison between different PBS buffers, the data analysis method from previous works^25,27,28^ was updated to remove the need for a clearly defined relative maximum. Both methods provide similar results, with the new method having slightly lower LODs, but the agreement between the approaches confirms the benefit of matching the baseline conductivity. In the following section, the sensor specificity is investigated by measuring the output voltages of the MERS-CoV spike, SARS-CoV-2 spike, and negative control (pure PBS buffer). The responses of the MERS-CoV spike proteins mimic the negative controls and demonstrate that the impedance sensor is capable of differentiating the target proteins and non-related samples. These experiments successfully verify the feasibility of using this nanowell sensor as a strong candidate to detect SARS-CoV-2 in human samples.

## Acknowledgement

Research reported in this publication was supported by the National Heart, Lung, And Blood Institute of the National Institutes of Health under Award Number U01HL150852. The content is solely the responsibility of the authors and does not necessarily represent the official views of the National Institutes of Health. Zhuolun Meng was supported by NSF grants 1846740 and 2002511.

