## Supplemental figures and table 1 for "A Label-free Nanowell-based Impedance Sensor for Ten-minute SARS-CoV-2 Detection"

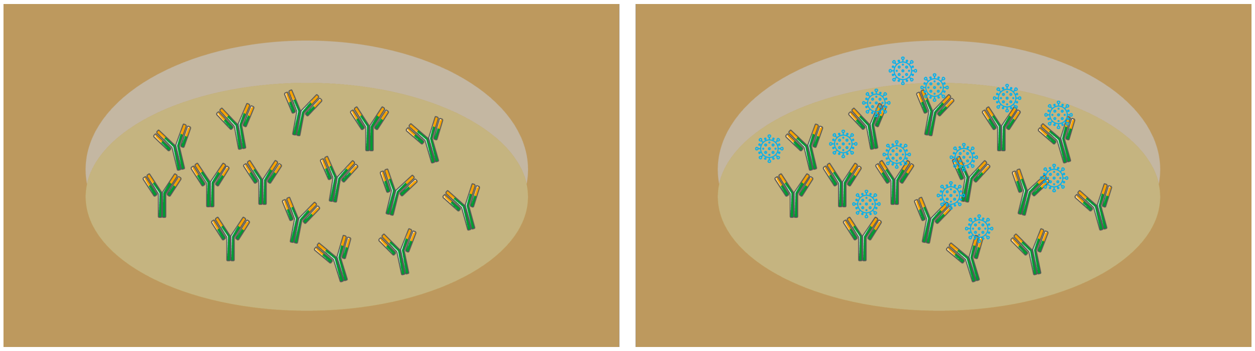

Figure S1: Top view of a single nanowell adsorbing antibodies and adsorbing target proteins

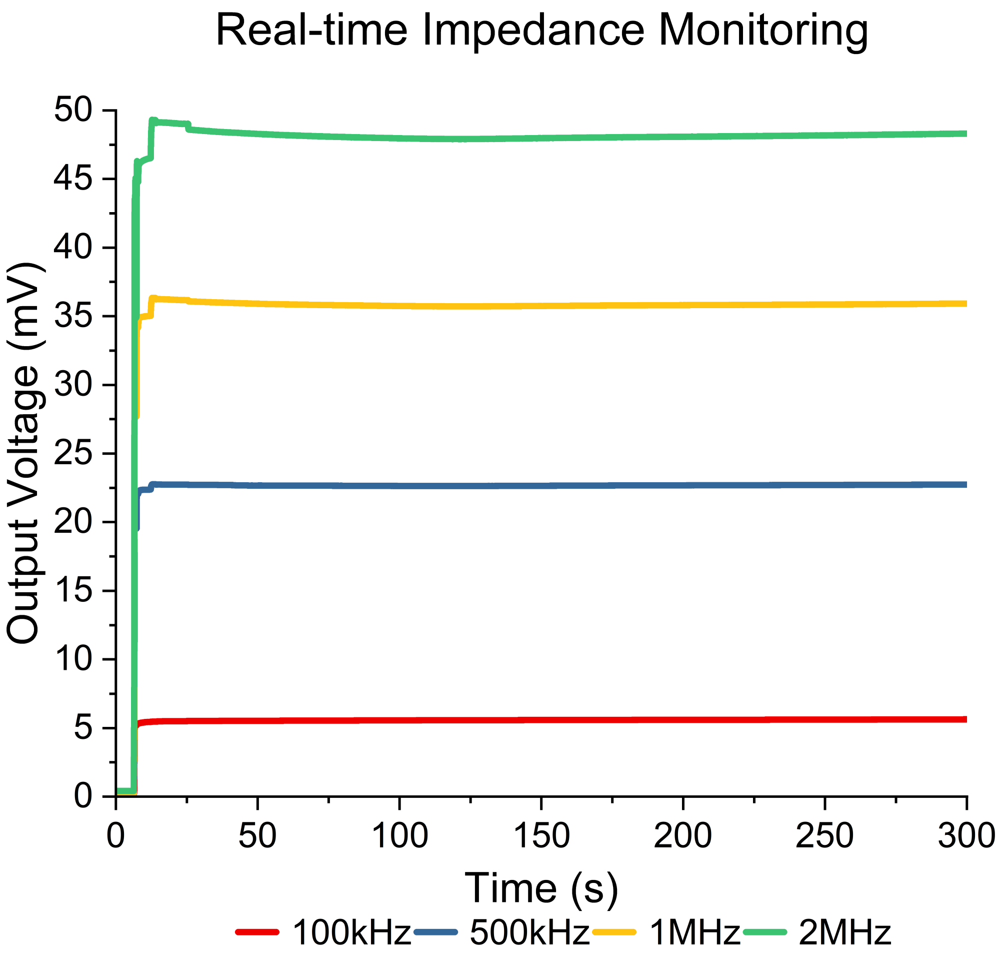


Figure S2: Real-time impedance monitoring four different frequencies under the liquid environment of 1X PBS

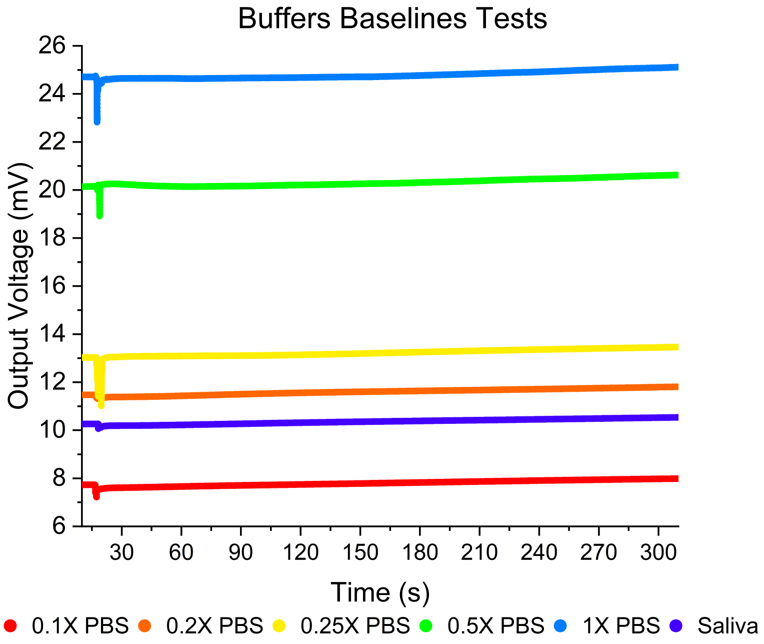


(a)

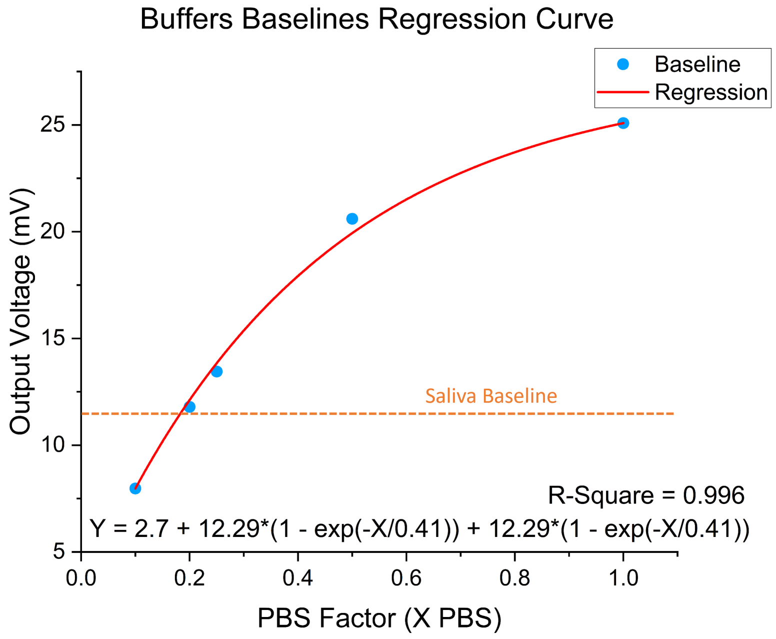


(b)

Figure S3: (a) Output voltage baselines for different concentrations of PBS and saliva. (b) Regression curve for different concentrations of PBS. Multiple experiments with different concentrations of PBS and saliva were performed, and the corresponding PBS concentration of saliva from the regression model is 0.18X PBS.

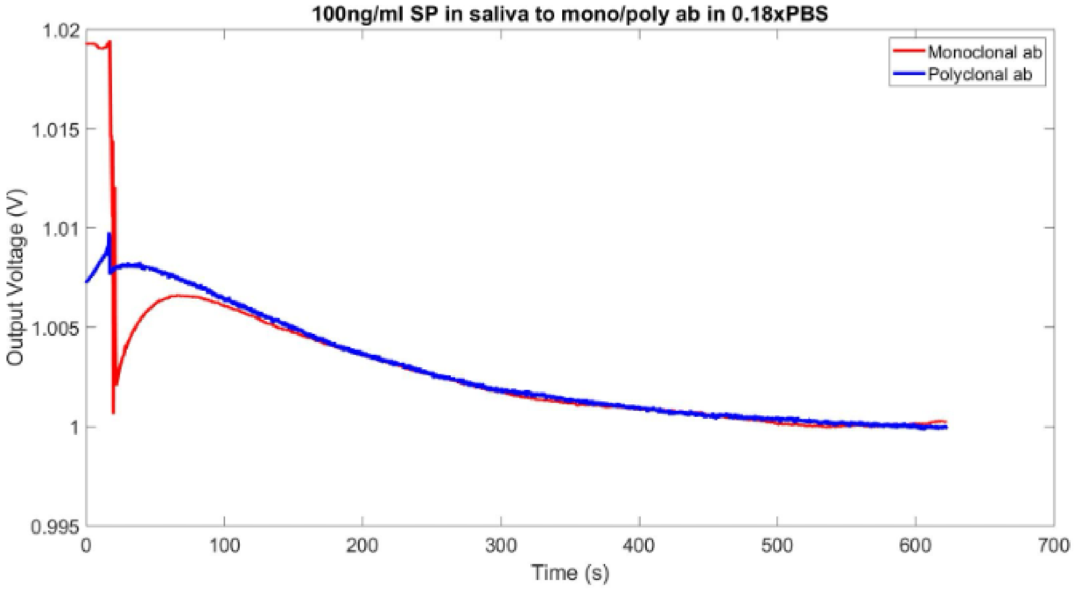


Figure S4: Comparison between Monoclonal and Polyclonal antibodies. The two curves have similar shapes and overlap around 150 seconds, which demonstrates that there appears to be no difference between Monoclonal and Polyclonal antibodies.

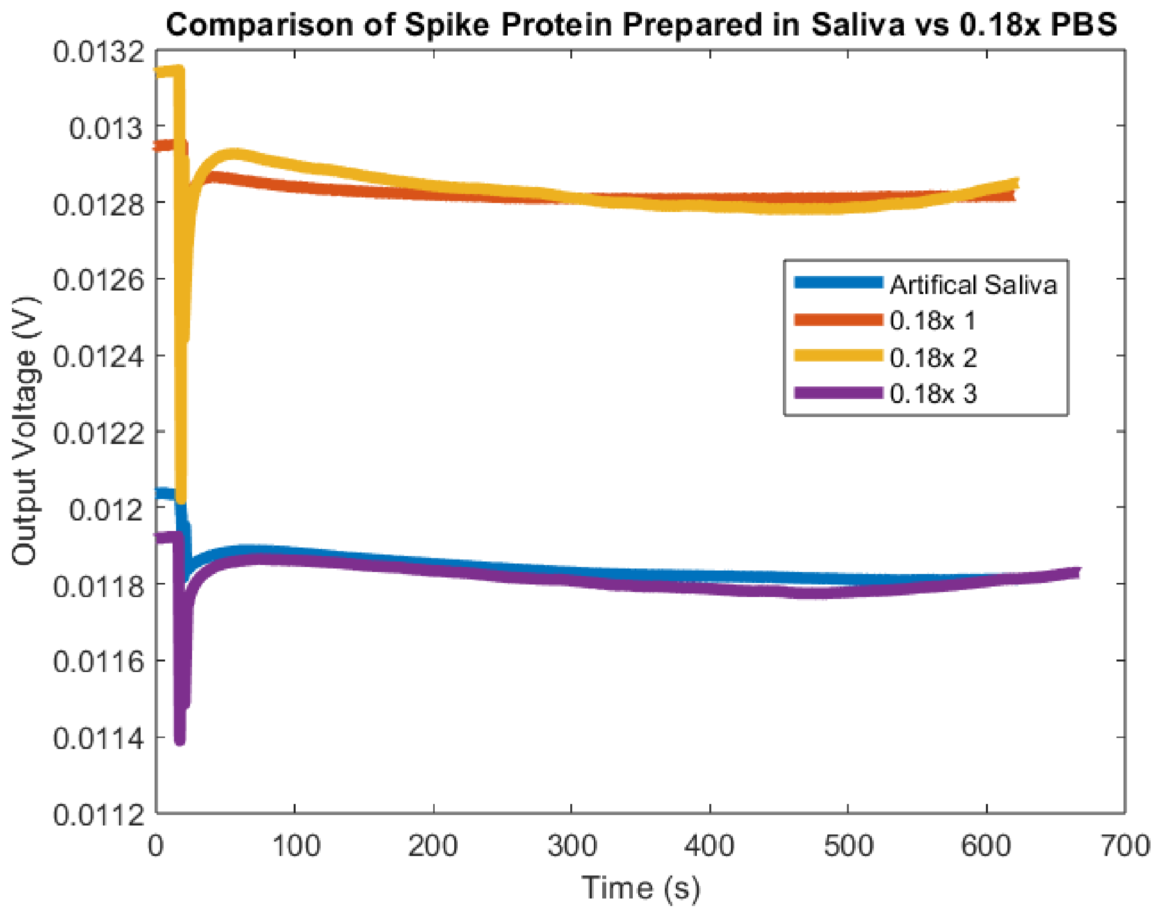


Figure S5: Comparison between SARS-CoV-2 spike protein antigen prepared in artificial saliva and 0.18X PBS. The antigen prepared in artificial saliva, shown in blue, has a similar output to the antigens prepared in 0.18X PBS. The voltage drop for spike protein prepared in artificial saliva is 0.49%, which is within the error bars for 100 ng/mL of spike protein prepared in 0.18X PBS.

Table S1: Table of detection limits for new and old data analysis methods using spike protein suspended in artificial saliva and antibodies suspended in 0.18X PBS for confidence levels ranging from 0.0001 - 0.05.

| **Analysis Method** | **Limit of Detection (ng/mL)** | **Limit of Detection (pM)** | **P** |
|---|---|---|---|
| New | 0.2 | 1.5 | $\leq 0.05$ |
| - | 0.5 | 3.7 | $\leq 0.001$ |
| - | 1 | 7.5 | $\leq 0.0001$ |
| Old | 1 | 7.5 | $\leq 0.0001$ |
